# Evidence for active regulation of transpiration in non-stomatal plants

**DOI:** 10.1101/2025.04.01.646543

**Authors:** Alicia V. Perera-Castro, Diego A. Márquez, Florian A. Busch, David Hanson

## Abstract

- Bryophyta (mosses) are a basal group of plants that lack stomata in their haploid form, as well as developed vascular tissue and a hydrophobic cuticle. Consequently, these plants are classified as poikilohydric, meaning poor control over water loss and are often assumed to reach equilibrium with ambient humidity. This classification does not fully align with the diverse strategies observed in mosses.
- We studied gas exchange in 14 species from Albuquerque and Boston, USA, under controlled dehydration conditions.
- Our results revealed significant variation in transpiration rates, cell wall equilibrium humidity, and desiccation times across species. These differences could not be explained by tissue water storage relative to the transpiring surface area, suggesting that water loss is not entirely passive. Additionally, species with better water control also presented traits of an avoidance strategy, including elastic tissues, high capacitance, and less negative osmotic potential, suggesting an adaptive constraint.
- These findings point to a basal, non-stomatal mechanism of water loss control through cell membranes and/or cell walls. Potentially, this mechanism is homologous to the non-stomatal control recently identified in angiosperms, which induces unsaturated conditions in the substomatal cavities. Bryophyta presents a valuable non-stomatal model for further investigating this mechanism and its evolutionary significance.

## INTRODUCTION

Poikilohydry refers to plants whose water status is completely dependent on their environment (Walter, 1931), i.e. inability to control water loss and isolate their cell water potential from that of the surrounding atmosphere. Poikilohydry is exhibited by lichens, gametophytes of ferns, lycophytes and bryophytes and some sporophytes of ferns (filmy ferns, Hymenophyllacea, Proctor, 2012). The shared characteristic of these poikilohydric group of plants is the absence of stomata. In bryophytes (mosses, liverworts and hornworts), stomata, when present, are limited to sporangia of the sporophyte and their physiological and evolutionary constraints differ from the stomata of leaves of tracheophytes (Renzaglia *et al*., 2020). Except for the very well-preserved gametophytes from the Lower Devonian (Pragian) Rhynie chert (Kenric & Crane, 1997; Edwards *et al*., 1998), no other fossils or extant gametophytes are known to have stomata (Duckett & Pressel, 2017). In contrast, homoiohydric plants maintain their water status within tight limits thanks to a more or less controlled stomata closure. This control ranges from active stomatal control in angiosperms to passive control in ferns and sporophyte of mosses and hornworts (Duckett & Pressel, 2022). Additional adaptations, such as hydrophobic cuticle, endohydric conducting apparatus, intracellular gas space system, and structures that can take up liquid water from the soil, further enhance their ability to avoid dehydration.

Such diversity in traits of different phylogenetic groups and life phases have made several authors consider a continuum poikilohydry-homoiohydry strategy in terrestrial plants (Raven, 2002; Proctor & Tuba, 2002; Raven & Edwards, 2004; Vitt *et al*., 2014). Some bryophytes share characteristics more associated with homoiohydric plants, such as the cuticle of some liverworts (Raven, 1977, 1984, 1993), endohydry of *Polytrichum* and filmy ferns (Hébant, 1977; Ligrone *et al*., 2000; Brodribb *et al*., 2020), and intracellular air-filled spaces of ventilated thalloid liverworts and hornworts (Duckett & Pressel, 2017). Furthermore, even when water conduction is typically external and diffuse, the growth habit of some bryophytes (gametophytes hereafter) resembles compacted shoots that confer canopies the ability to retain large amounts of external capillary water (Dilks & Proctor, 1979; Proctor *et al*., 1998). This also results in thick boundary layers, which allow bryophytes to lose this external water slowly without affecting cell water status (Marcshall & Proctor, 2004; Rice *et al*., 2014). These characteristics, therefore, buffer the rate of water loss for a given availability of water in the environment and delay water stress (Proctor *et al*., 2007). Such “avoidance” behavior of moss gametophytes has been rarely considered when discussing the poikilohydry-homoiohydry continuum (Vitt *et al*., 2014; Jabłońska *et al*., 2023), even though this short-term storage of external capillary water is not incompatible with the ability to tolerate desiccation (up to -100 MPa; Oliver *et al*., 2020), a trait also widely observed among the studied bryophytes (Morales-Sánchez *et al*., 2022).

In addition to the different levels of poikilohydry that can be defined according to the buffering effect of a canopy structure, bryophytes may present also differences in their shoot-level water storage. In addition to their high water content of the cytoplasm and capacitance of bryophytes (Proctor & Tuba, 2002, Perera-Castro *et al*., 2020a), bryophytes present the thickest cell wall of terrestrial plants (0.6-3.5 µm, Perera-Castro *et al*., 2022a), which comprises the 4%-18% of water content of the tissues (Proctor *et al*., 1998; Proctor, 1999; Perera-Castro & Flexas, 2022). Cell wall thickness has been positively correlated with the carbon uptake of mosses in a controlled dehydration treatment at 33% of relative humidity (Coe *et al*., 2019), suggesting that cell walls themselves act as a reservoir for water. However, neither the canopy nor the shoot buffering effect explained above can be considered an active water control, since it does not affect the water transport conductance of tissues during dehydration but rather the amount of water lost before impacting water potential.

Among the components involved in the water pathway to the sites of evaporation (cytoplasm – cell plasma membranes – cell wall), only the plasma membranes and its structures can serve as a true barrier controlling water loss, for example, through the widely studied aquaporins (AQPs). Those ubiquitous transmembrane proteins act as channels for many molecules, above all water, and have been described to be involved in a wide variety of processes, including water fluxes through plant tissues that are linked to the transpiration stream (Van Dongen & Borstlap, 2004; Maurel *et al*., 2015; Kapilan *et al*., 2018; Singh *et al*., 2020; Byrt *et al*., 2023). Consistent with this observation, different AQP expressions resulted in different water control and drought tolerance in tracheophytes (Kapilan *et al*., 2018; Verdoucq & Maurel, 2018; Singh *et al*., 2020; Byrt *et al*., 2023). The study of AQPs in the moss *Physcomitrella patens* has revealed that this species contains the same subfamilies of AQPs that are described in angiosperms (PIP, TIP, NIP and SIP subfamilies), plus other exclusive subfamilies (XIP, HIP, GIP). This indicates that the main radiation of plant AQPs preceded the land plant colonization (Borstlap, 2002; Danielson & Johanson, 2008). In addition, Liénard *et al*. (2008) reported that the suppression of AQPs in *Physcomitrella* could increase its sensitivity to moderate stress conditions (visually evaluated by the shrinkage of shoots). However, the authors stated that “[although] AQPs are not required on the cell surface in contact with the air, AQPs could delay cellular water loss by facilitating water exchanges through the part of the cell surface in contact with the liquid phase”. That is, AQPs are presented as facilitators of water entrance rather than as controllers of water loss. Since their description in mosses, aquaporins have emerged as important components of the transcriptional response to water stress in this group of plants (Cuming *et al*., 2007), although researchers have been reticent to consider this as a way of water control: “… their role in mosses has been unclear, as poikilohydric plants do not regulate their water potential” (Charron & Quatrano, 2009).

We argue that such discoveries serve as a starting point for considering bryophyte gametophytes as poikilohydric, non-stomatal plants with a certain level of water control at the shoot level. The regulation of water flux through cell membranes by AQPs has been proposed as a non-stomatal mechanism controlling transpiration in leaves (Wong *et al*., 2022), challenging the long-standing assumption that leaf air spaces are nearly saturated with water vapor (Diao *et al*., 2024). It has been reported that relative humidity in these spaces can drop to 80% under increasing vapor pressure deficit, while mesophyll cells maintain turgor (Cernusak *et al*., 2024). Even further, this humidity can drop as low as 60% when using transgenic lines with non-closing stomata (Cernusak et al., 2019). Leaves with non-closing stomata can be considered nearly “cell-wall-naked” to atmospheric conditions, similar to those found in poikilohydric plants. However, the epidermal cells and cuticle of the leaf still provide an additional layer of isolation for the mesophyll cells, influencing how non-stomatal control of transpiration is studied and its physiological implications. Notably, if bryophytes —which lack stomata in the gametophyte— exhibit this form of transpiration control, it can not only be studied more directly but also provide insights into its ancient evolutionary origins. As with the lack of saturation in the leaf air space, low relative humidities on the surfaces of bryophytes— without a corresponding reduction in cytosolic water potential—would represent a non-stomatal water control mechanism through plasma membranes. If this mechanism is confirmed in mosses, it would further support the idea of water control in poikilohydric organisms and suggest a more primitive origin for the non-stomatal control of transpiration.

The aim of this study was to investigate water control in the gametophytes of mosses by measuring gas exchange in detached moss shoots during dehydration. We used 14 species collected from Albuquerque and Boston. They were maintained in a growth chamber at the University of New Mexico. Pressure-volume traits and microscopy analysis were performed to gain further insight into moss water relations and strategies. These analyses included measurements of apoplastic and cytosolic water potential, capacitance, osmotic potential, and cell wall thickness. Our hypothesis was that the cell wall water potential of mosses would undergo a similar drop to that reported for angiosperms mesophyll cells, also without a loss of turgor or equilibration with cytosolic water potential —i.e. mosses exhibit a non-stomatal control of water loss as well.

## MATERIAL AND METHODS

### Plant material

A total of 14 species of mosses (Bryophyta phylum) were collected in the field in Sandia Crest (Albuquerque) and Billerica (Boston), in the US (Table 1). Patches of 1-5 cm^2^ of moss and below-moss soil/wood were introduced in non-sealed plastic bags and stored in a growth chamber at the University of New Mexico. The moss patches were placed together in non-hermetically sealed plastic boxes covered with a transparent plastic film. All plastic boxes were randomly distributed inside the growing chamber. Mosses were maintained irradiated with 430-460 µmol m^-2^ s^-1^ of light with a photoperiod of 14/10 h and a moss temperature of 21-23°C during the day and 16-20 °C during the night. Moss temperatures were measured by placing a thermocouple on the canopy surface of the mosses (EL-USB-TC, Lascar Electronics Inc, USA). Mosses’ surfaces were sprayed with distilled water once a week. All measurements were performed 1-3 months after collection. Species were identified according to Allred *et al*. (2024) key.

**Table 1.** List of studied species.

| Species | Family | Location | Growth habit |
| --- | --- | --- | --- |
| *Atrichum undulatum* (Hedw.) P. Beauv. | Polytrichaceae | Billerica, MA | Acrocarpous |
| *Ceratodon purpureus* (Hedw.) Brid. | Ditrichaceae | Sandia Crest, NM | Acrocarpous |
| *Dicranium* sp Hedw. | Dicranaceae | Billerica, MA | Acrocarpous |
| *Hygrohypnum eugyrium* (Schimp.) Loeske | Amblystegiaceae | Billerica, MA | Pleurocarpous |
| *Hypnum cupressiforme* Hedw. | Hypnaceae | Sandia Crest, NM | Pleurocarpous |
| *Hypnum pallescens* (Hedw.) P.Beauv. | Hypnaceae | Billerica, MA | Pleurocarpous |
| *Leucobryum albidum* (Brid. ex P.Beauv.) Lindb. | Leucobryaceae | Billerica, MA | Acrocarpous |
| *Plagiomnium cuspidatum* (Hedw.) T. Kop. | Mniaceae | Billerica, MA | Acrocarpous |
| *Pohlia nutans* (Hedw.) Lindb | Mniaceae | Sandia Crest, NM | Acrocarpous |
| *Polytrichum commune* Hedw. | Polytrichaceae | Billerica, MA | Acrocarpous |
| *Thuidium delicatulum* (Hedw.) Schimp. | Thuidiaceae | Billerica, MA | Pleurocarpous |
| *Trichostomopsis umbrosa* (Müll.Hal.) H.Rob. | Pottiaceae | Sandia Crest, NM | Acrocarpous |
| Acrocarpic: compact growth with capsules at the tip of the stems |  |  |  |
| Pleurocarpic: spreading growth with capsules along the sides of the stems |  |  |  |

### Gas exchange measurements

Measurements of net CO_2_ assimilation rate (*A*_N_) and transpiration rate (*E*) in response to variations in water content (‘dehydration curve’) were done with a LI-6800 gas exchange system (LiCOR Biosciences, Lincoln, NE, USA). Samples were saturated to excess with distilled water prior to the measurements. Then, they were placed over a custom-made cuvette consisting of a 6 cm^2^ gasket fixed to a piece of thin polyester stocking fabric as detailed in Perera-Castro *et al*. (2020b). The water concentration of the incoming air (reference water) was set to 10 mmol mol^−1^ air. The flow rate within the chamber was maintained at 400 μmol s^−1^ and the CO_2_ concentration in the incoming air was set at 420 μmol mol^−1^ air. Irradiance was set at a saturation level of 800 μmol m^−2^ s^−1^ standardized for all species (tested previously with fast light-response curves). The temperature inside the chamber was maintained at 26.6±0.5 °C. Gas exchange recordings were alternated with weighing of the sample each 2-5 min to obtain the fresh weight (FW) during the dehydration curve. Water content (WC) at any given time during dehydration was calculated as: WC=(FW-DW)/DW, where DW is the dry weight obtained at the end of the experiment by keeping the samples at 70°C for 48h till constant weight. An entire cycle of desiccation lasted for 20-45 min and stopped when *A*_N_ was close to zero. Due to low gas exchange rates, measurements of CO_2_ leakages (Flexas *et al*., 2007) were done regularly with an empty cuvette at the beginning and end of each dehydration curve. *A*_N_ and *E* at optimum WC (WC_opt_, i.e., at maximum *A*_N_) were termed *A*_opt_ and *E*_opt_, respectively. Maximum *E* (*E*_max_) along the entire dehydration curve could be observed at optimum WC or higher. *A*_N_ and *E* were expressed per projected shoot area in both acrocarpic and pleurocarpic species, obtained by an image processing program (ImageJ, NIH, Rockville, MD, USA). Shoot mass area (SMA) was calculated as the ratio of DW and the projected shoot area. For each species, a total of 3-5 dehydration curves were performed.

### Pressure-volume curves

Between 4 and 6 pressure–volume curves per species were performed (except for *Dicranium* sp, for which *n*=1 due to the scarcity of plant material) by slowly air-drying full hydrated samples and alternately weighing and measuring its water potential (Ψ_w_) by using a psychrometer (model WP4C, Decagon Device Inc.). The turgid weight (TW) of each specimen was estimated as the x-intercept of Ψ_w_ versus fresh weight at any time on the curve (FW), avoiding the “plateau effect” of extracellular water (Proctor *et al*., 1998; Hájek & Beckett, 2008). Both water content (WC) and relative water content (RWC) were calculated, the latter as RWC=100(FW-DW)/(TW-DW). The saturating water content (SWC) was calculated as SWC=(TW-DW)/DW and SWC_area_=(TW-DW)/projected shoot area. As for gas exchange samples, the projected shoot area was obtained with ImageJ and the shoot mass area (SMA) was also calculated as the ratio of DW and the projected shoot area. The point from which the pressure-volume curve (-1/Ψ_w_ vs RWC) became linear was considered the turgor loss point. RWC, WC and Ψ_w_ measured at that point were obtained (RWC_tlp,_ WC_tlp_ and π_tlp_, respectively). Osmotic potential at full rehydration (π_o_) was calculated as the inverse of the y-intercept of the linear regression of the pressure–volume curve; this is the linear portion under the turgor loss point. The bulk modulus of elasticity (ε) was calculated as the slope of pressure potential versus total RWC. Absolute capacitance at full turgor (*C*_ft_) and at turgor loss point (*C*_tlp_) was determined from the relationship between Ψ_w_ and RWC above and below turgor loss point, respectively. The extracellular apoplastic fraction (*a*_f_) was considered the fraction of RWC when osmotic potential approaches-∞ and was calculated as the x-intercept of the pressure– volume curve. The equivalent WC that corresponds to *a*_f_ (WC_apo_) was also obtained for the - 1/Ψ_w_ – WC relationship. Cytosolic (or symplastic) fraction (*c*_f_) and cytosolic water content (WC_cyt_) were also obtained as *c*_f_=1-*a*_f_ and WC_cyt_=SWC-WC_af_.

### Microscopy and anatomy analysis

Three shoots per species were selected for microscopy analysis. Sample manipulation and cross sections of leaves (phyllids) were manually done under a stereoscopic microscope. Images of cross sections and leaf surfaces were obtained with a camera (CMOS USB 3.0 10MP Camera) attached to a light microscope (Leica DM6 B Microscope). Images taken with a magnification of 400X were analysed with ImageJ for obtaining the cell wall thickness (*T*_cw_) from the cross-section samples, and cell areas (*Cell*_area_) from leaf surface images. Each replicate (*n*=3) represents the average of 3 to 20 measurements that were taken for each parameter and sample whenever possible. As an indirect measurement of the total exposure surface area, measurements of the total leaf area per shoot area (*A*_leaf_/*A*_shoot_) were done by detaching all leaves of a shoot (*n*=1). A stereoscopic microscope was used in some species. For *Pleurozium schreberi* and *Thuidium delicatulum*, *A*_leaf_/*A*_shoot_ was assumed to be close to 1, due to the tiny size of their dispersed leaves.

### Calculation of cell wall water potential

At any moment of the dehydration curve, the water vapour concentration over the cell wall (*W*_cw_) was calculated from the Fick’s law of diffusion equation from a non-stomatal interrupted cell wall through the boundary layer to the mixed atmosphere:

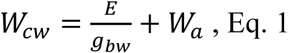

where *g*_bw_ is the boundary layer conductance for water and *W*_a_ is the atmosphere water vapor concentration (air coming out of the chamber). *g*_bw_ was obtained from LI-6800 empirical calculations for a broad surface, assuming *g*_bw_ is the same on both sides of the surface. Then, the saturated water vapor pressure over the cell wall (*W*_cw,sat_) was calculated as a function of shoot temperature (Buck, 1981):

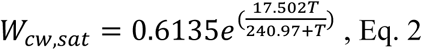

where *T* is the temperature of the shoot, estimated from energy balance calculations of LI-6800. The relative humidity at the cell wall surface (RH_cw_) was then calculated as:

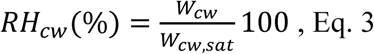

Since the shoot surface cannot be considered completely flat, impacting *g*_bw_, further affecting temperature estimates by energy balance that also depend on *g*_bw_, a sensitivity analysis was performed to determine the variability of RH_cw_ with changes in *g*_bw_ (Fig. S1) and *T* (Fig. S2). An absolute variation of *g*_bw_ in 1.6 mol m^-2^ s^-1^ did not provoke significant variation in RH_cw_ directly or by changes in energy balance *T* (Fig. S1). An averaged change in RH_cw_ of 5.1%±0.8% for 1°C of error in estimates of *T* was calculated for all species (Fig. S2). The sensitivity of energy balance *T* to the moss absorbance (α) was also estimated (Fig. S3), resulting in a maximum change in *T* of 0.5°C for a variation of α between 0.6 and 1.

Water vapor potential at the cell wall (Ψ_cw_) was calculated by the Kelvin equation as described in Nobel (2020):

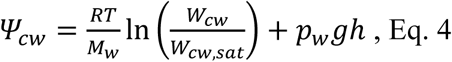

where *R* is the universal gas constant (8.314 J mol^−1^ K^−1^), *T* is the temperature, *M*_w_ is the partial molar volume of water (18 × 10^−6^ m^3^ mol^−1^), *p*_w_ is water density, *g* is the gravitational acceleration and *h* is altitude (1508 m a.s.l.). The equation was conveniently rescaled to MPa (*RT*/*M*_w_ = 137.3 MPa).

Since most of the water of a non-fully hydrated shoot is located within the living cells, this likely dominates the determination of whole-tissue water potential, as assumed for leaves (Scoffoni *et al*., 2023). Therefore, cytosolic water potential (Ψ_cyt_) was assumed to be equal to Ψ_w_ estimated at any moment of the dehydration curve from the -1/Ψ_w_ - WC relationship of the pressure-volume curves (Fig. S4).

### Statistical analysis

All analyses were performed using the R statistical software (R Core Team, 2023). Some special packets were used: *plyr* (Wickham, 2011), *ggplot2* (Wickham, 2016). Intraspecific differences in transpiration rate at different moments of dehydration were evaluated by a *t*-test. Differences in the transpiration rates of the studied species were evaluated by the Kruskal-Wallis test. In order to discern the biological/methodological origin of such differences in transpiration, linear regression between transpiration and related chamber conditions (vapor pressure and temperature) was performed. Linear regression was also used for testing the relationship between the calculated RH_cw_ and possible determinants, like boundary layer conductance involved in its calculation, or morphological parameters theoretically involved with the amount of water available in the sample for its exchange, such as saturated water content per shoot area or the exchange area of samples. The relationship between RH_cw_ and parameters derived from the pressure-volume was also explored.

## RESULTS

### Mosses present different transpiration rates and dehydration kinetics

*A*_N_ responses to dehydration described a parabolic curve in all studied species, with a decrease in *A*_N_ for any WC higher or lower than WC_opt_, indicating that both excess of hydration and dehydration can be detrimental for *A*_N_ (Fig. 1a). WC_opt_ varied between 1.9±0.1 g H_2_O g^-1^ DW in *Polytrichum commune* to 6.3±0.3 g H_2_O g^-1^ in *Leucobryum albidum*. By contrast, the transpiration rate remained at its maximum value (*E*_max_) for any WC higher than WC_opt_ (Fig. 1b). This asymptotic value of *E*_max_ was significantly higher than *E*_opt_, except in *Pohlia nutans* and *Thuidium delicatum* (*t*-test, *P*>0.001), indicating that in most studied species, the transpiration rate started decreasing before maximum assimilation rate during dehydration. Significant differences between species were observed in both *E*_opt_ (Fig. 2a) and *E*_max_ between species, ranging from a *E*_opt_ of 30.0±0.5 µmol m^-2^ s^-1^ in *Polytrichum commune* (*E*_max_ = 43.3±0.9 µmol m^-2^ s^-1^) to a *E*_opt_ of 12.9±2.0 µmol m^-2^ s^-1^ (*E*_max_ = 14.9±2.9 µmol m^-2^ s^-1^) in *Pohlia nutans*. Such differences were not correlated with *A*_opt_ (data not shown) or with the vapor pressure inside the chamber (Fig. 2b), which resulted from an air temperature interspecific variation of 26.6 ± 0.5°C and an incoming H_2_O concentration of 9.98 ± 0.03 mmol mol^-1^. The time required for dehydration (from *A*_opt_ to *A*_N_ = 0), which took from 8 to 31 minutes, was not significantly correlated with *A*_opt_, *E*_opt_ or *E*_max_.

**Figure 1.**
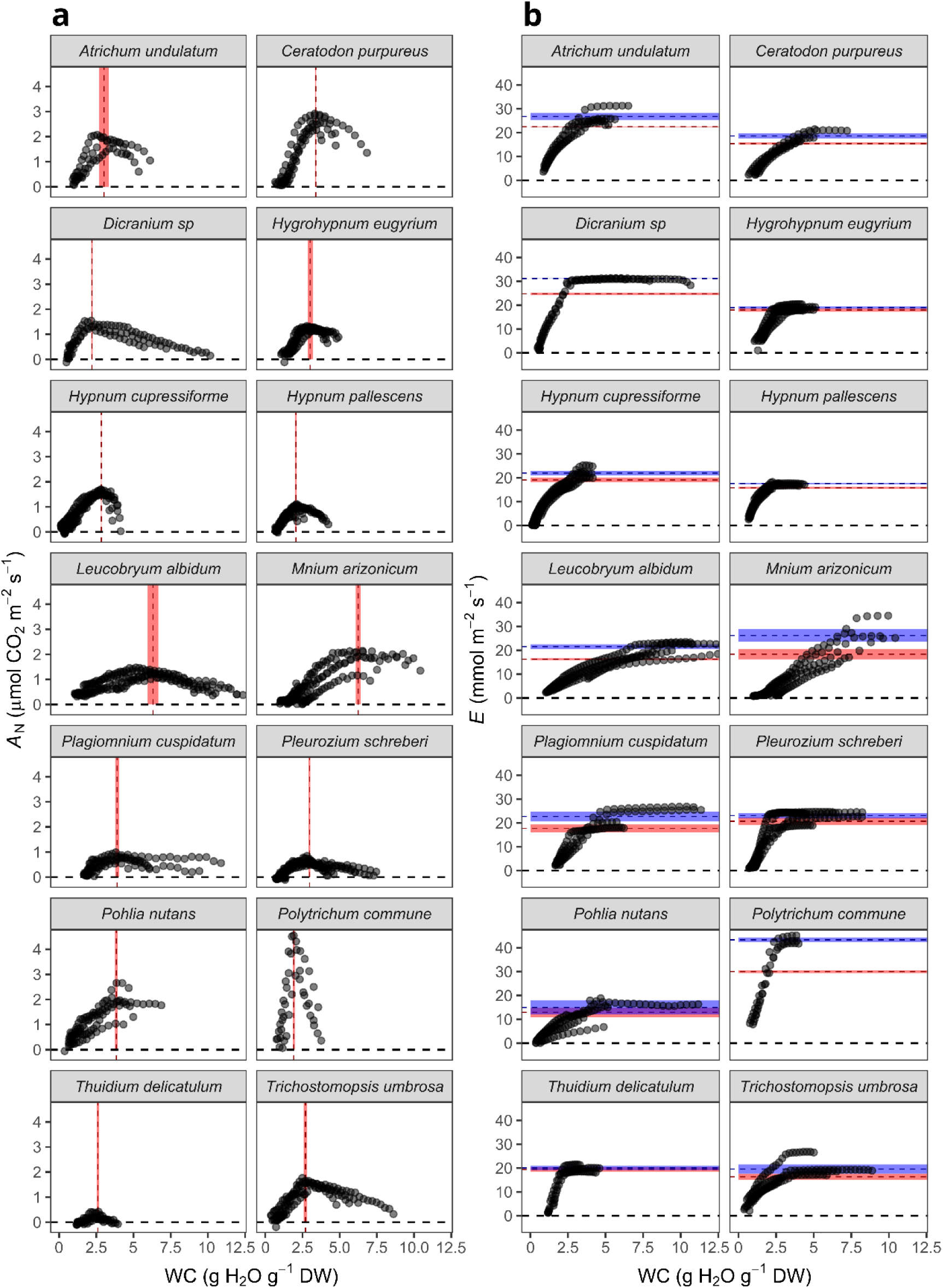
Response to dehydration of (a) net CO_2_ assimilation rate (*A*_N_) and (b) transpiration rate (*E*) for the studied species. Red vertical lines indicate the average water content (WC) at which maximum *A*_N_ (*A*_opt_) is found (WC_opt_). Red and blue horizontal lines indicate averaged transpiration rates at WC_opt_ (*E*_opt_) and maximum transpiration rates (*E*_max_), respectively. Shaded areas indicate ±standard errors of the respective parameters.

**Figure 2.**
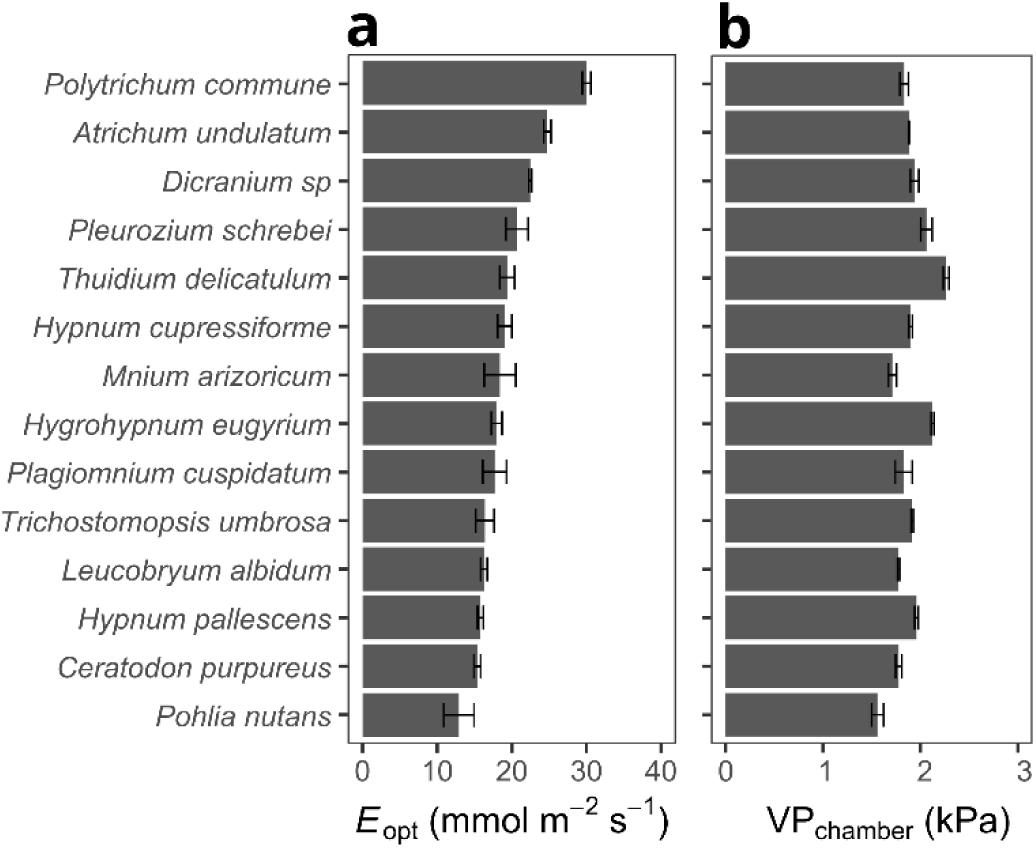
Interspecific variation of the (a) transpiration rate at the maximum net CO_2_ assimilation (*E*_opt_) and (b) vapor pressure inside the chamber (VP_chamber_) at an air temperature of 26.6 ± 0.5°C and an incoming H_2_O concentration of 9.98 ± 0.03 mmol mol^-1^. Both *E*_opt_ and VP_chamber_ significantly differ between species (*P*<0.001, Kruskal-Wallis test), although associated (non-significant linear regression).

### Moss cell walls can sustain lower water potential at maximum *A*_N_ without cytosolic dehydration

At WC_opt_, the calculated values of RH_cw_ showed significant differences between species, from 100.8%±3.0% to 59.4%±4.7% in *Polytrichum commune* and *Pohila nutans*, respectively, which corresponded to values of Ψ_cw_ from 0 to -58.9 MPa (Fig. S5). The parameter *E*_opt_ explains most strongly the variations in RH_cw_ observed between species (*R*^2^=0.585, *P*=0.001, Fig. 3a). No significant correlation exists between RH_cw_ and the *g*_bw_ at this WC_opt_ (Fig. 3b). RH_cw_ is not explained by the amount of water that samples can provide per unit area, as evidenced by the lack of correlation between RH_cw_ and SWC_area_ (Fig. 3c), both in the apoplastic and the symplastic (Fig. S6). No anatomical parameter examined here can explain differences between RH_cw_ among species, including those that alter the exchange surface, such as the *A*_leaf_/*A*_shoot_ ratio or the *A*_cw_ (Fig. 3d).

**Figure 3.**
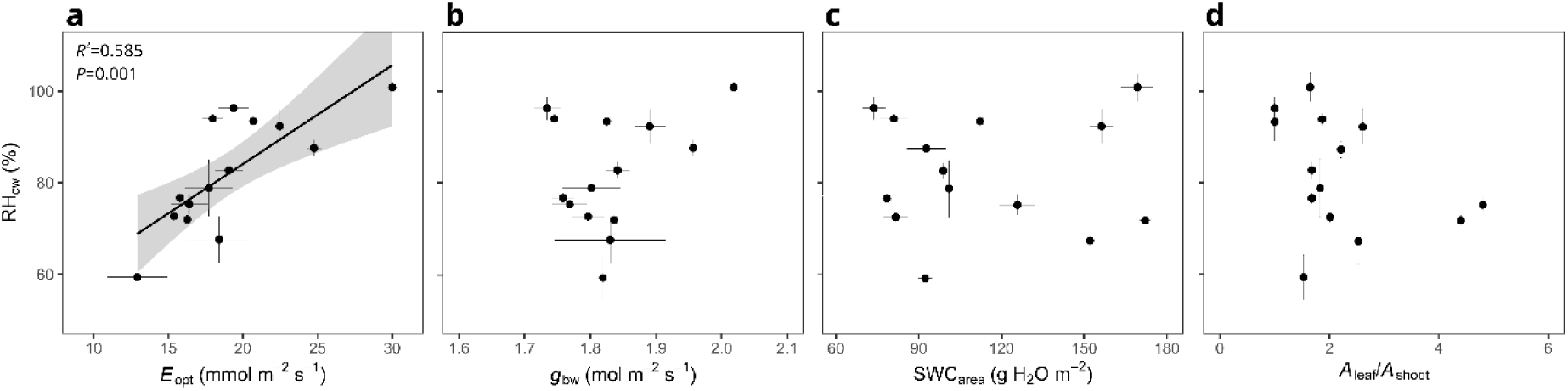
Relationship between relative humidity at the cell wall (RH_cw_) at optimum water content (where the assimilation rate is maximum) and (a) transpiration rate at optimum water content (*E*_opt_), (b) conductance of the boundary layer to the diffusion of water (*g*_bw_), (c) saturated water content per unit shoot area (SWC_area_), and (d) ratio of leaf area per shoot area (*A*_leaf_/*A*_shoot_). Points represent the mean ±SE for each species (n=3-5).

For WC<WC_opt_, Ψ_cw_ was significantly lower than Ψ_cyt_ in all studied species, so that when *A*_N_ was still positive in most of the species and Ψ_cyt_ was equal to π_tlp_, Ψ_cw_ could be 10 orders of magnitude more negative than Ψ_cyt_ (Ψ_cw_/Ψ_cyt_ at turgor loss point was between 78.8±4.7 for *Leucobryum albidum* and 23.1±1.9 for *Polytrichum commune*) (Fig. 4). Four out of five dehydration curves of *Hygrohypnum eugyrium* reached *A*_N_ = 0 (and the dehydration curve was interrupted) before reaching the turgor loss point. For all other species, *A*_N_ was still positive when the plants crossed π_tlp_, indicating these plants are still active at low cell wall water potential.

**Figure 4.**
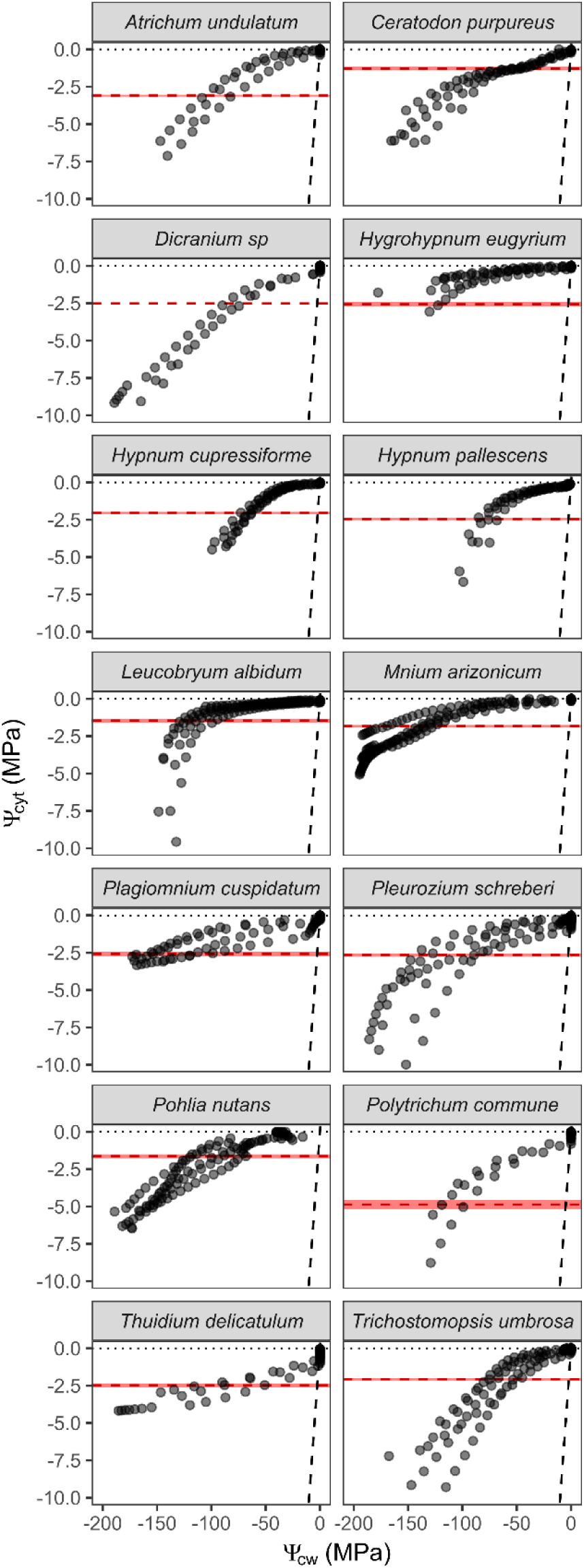
Relationship between cytoplasmic water potential (Ψ_cyt_) and cell wall water vapor potential (Ψ_cw_) for the dehydration curves of the studied species. Ψ_cyt_ was calculated from the WC at any moment of the dehydration curve by using the relationship between -1/Ψ_w_ and WC of pressure volume curves (Fig. S4) and assuming Ψ_w_=Ψ_cyt_. The horizontal dashed red line represents the averaged water potential at the turgor loss point (π_tlp_). The diagonal dashed black line indicates a 1:1 relationship.

### Cell wall surface humidity correlated with dehydration avoidance traits

Some pressure-volume derived parameters, which were measured in independent samples of the same species, showed a significant correlation with the relative humidity calculated for cell walls (Fig. 5a,b,c). Highest RH_cw_ values were measured in species with the highest bulk modulus of elasticity (higher ɛ, more rigidity, Fig. 5a), with the lowest capacitance at full turgor (*C*_FT_, Fig. 5b) and with more negative osmotic potential (π_o_, Fig. 5c). The species with the highest cytosolic water content per unit dry weight (WC_cyt_) presented with the lowest RH_cw_ (Fig. 5d), independently of the apoplastic water content (Fig. 5e). These profiles of traits (more elastic tissues, higher capacitance, low osmotic adjustment and high-water storage) corresponded to the species whose dehydration curves took longer to finalize (from *A*_opt_ to *A*_N_=0, Fig. 5f).

**Figure 5.**
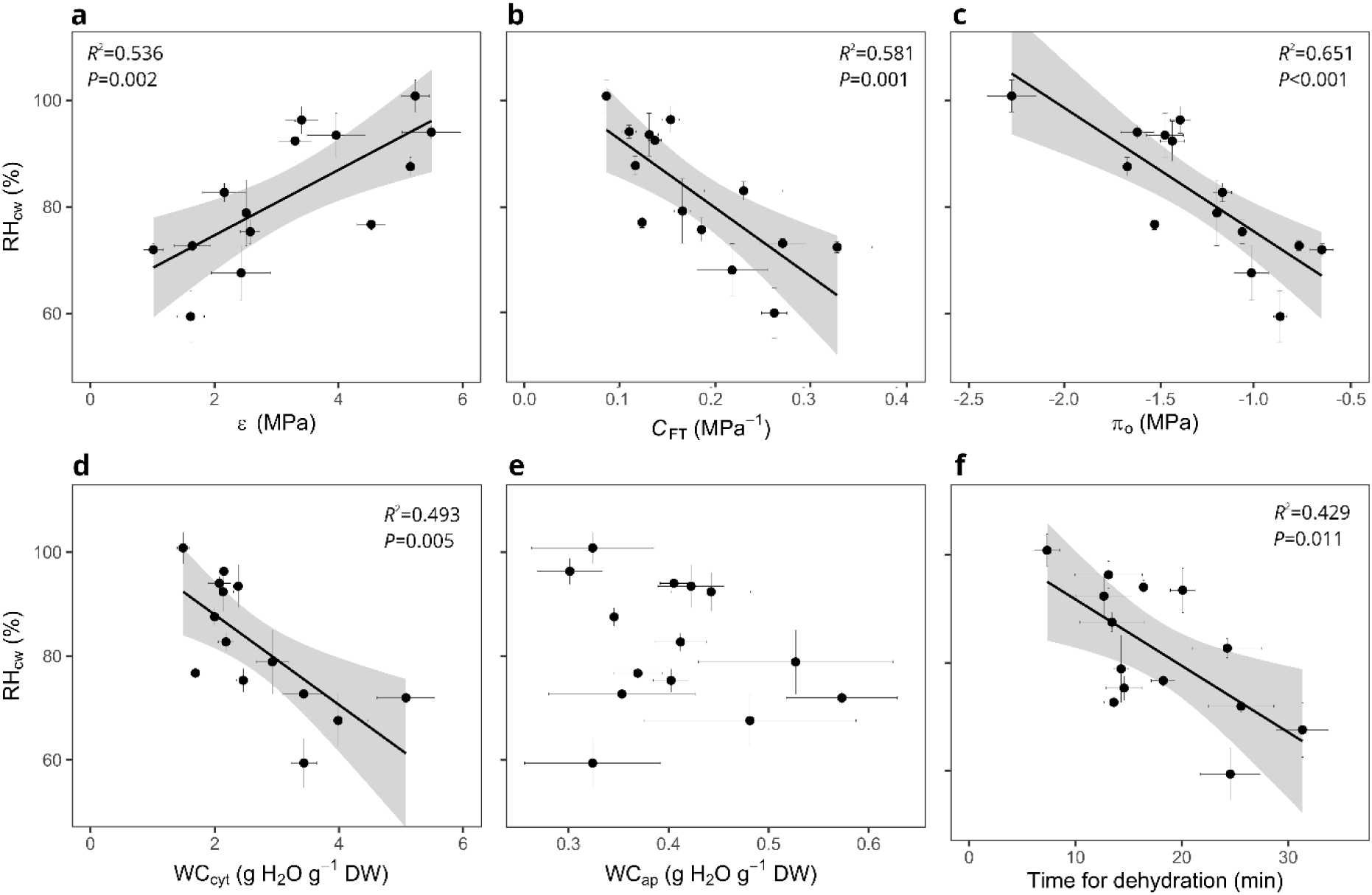
Relationship between relative humidity at the cell wall (RH_cw_) and water relation related traits: (a) bulk modulus of elasticity (ɛ), (b) capacitance for water content range above turgor loss point (*C*_FT_), (c) osmotic potential (π_o_), (d) cytosolic water content per unit dry weight (WC_cyt_), (e) apoplastic water content per unit dry weight (WC_ap_), and (f) time for dehydration (from *A*_opt_ to *A*_N_=0 in the dehydration curves). Points represent the mean ±SE for each species (*n*=3-5 for RH_cw_ and time for dehydration, *n*=1-6 for pressure-volume derived parameters).

## DISCUSSION

### Evidence for actively controlled water loss in mosses

As introduced, poikilohydry has been conceived as an inability to control water loss, equivalent to a nearly infinite conductance of tissues to water only driven by capillary forces and boundary layer conductance (Raven, 2002; Proctor & Tuba, 2002; Raven & Edwards, 2004; Vitt *et al*., 2014). In effect, the reported shoot maximum rates of transpiration for the studied mosses (15-26 mmol m^-2^ s^-1^) are more than 10-fold higher than those reported for homoiohydric plants (below 4 mmol m^-2^ s^-1^ for most reports, Kattge *et al*., 2020, TRY database, mostly angiosperms), whose gas exchange capacities are limited mainly by stomatal density and stomatal pore size under non water stress conditions (Franks *et al*., 2015; Fanourakis *et al*., 2015). However, from this study, three observations support the idea of only partially passive water control in mosses. First, although certainly high, the transpiration rates differed between the studied species of mosses under similar measuring conditions of vapor pressure of the chamber and dryness of the incoming air. This result was unexpected for detached and non-overlapping shoots that do not conserve their canopy water retention properties. Second, during dehydration of excessively hydrated cells, transpiration rates start decreasing before registering the maximum assimilation rate (*A*_opt_), i.e., when WC is still higher than WC_opt_. This is evidenced by the general significant differences between *E*_max_ and *E*_opt_ (Fig. 1). Typically, *A*_opt_ of bryophytes is considered close to 100% RWC, when there is no external water that impedes CO_2_ diffusion or cell dehydration that affects the biochemistry of photosynthesis (Dillks & Proctor, 1979; Deltoro *et al*., 1998; Rice *et al*., 2011; Wagner *et al*., 2013; Perera-Castro *et al*., 2020ab, 2022b). Thus, the transpiration rate apparently started decreasing before dehydration had an effect on the internal water content of cells. And third, at the moment when WC_opt_ was reached, humidity values at the surface of the cell wall (RH_cw_) calculated from *E*_opt_ were lower than 100% in most species, going as low as 60% in *Pohlia nutans*. As expected, RH_cw_ and the equivalent water potential (Ψ_cw_) decreased together with transpiration during desiccation. However, at the turgor loss point, Ψ_cw_ is clearly more negative than the water potential estimated from pressure-volume curves for the corresponding water content. A difference in water potential between the cytoplasm and the cell wall surface requires a finite hydraulic conductance across the plasma membrane, and the decoupled evolution of Ψ_cw_ and Ψ_cyt_ (Fig. 4) are consistent with principles of dynamic flow control.

The advanced response of transpiration to dehydration, the values of RH_cw_ lower than 100%, even at WC_opt_ where internal cellular dehydration is not expected, and the discrepancies between Ψ_cw_ and Ψ_cyt_ point to a water loss that is in part actively regulated and speaks against a fully passive water loss that is frequently assigned to mosses. The range of RH_cw_ values reported in our study (60-100%) for mosses at WC_opt_ are close to those measured for angiosperms internal airspaces (60-93%, Cernusak *et al*., 2024), including those studies where stomatal opening is induced somehow (60-70%, Jarvis & Slatyer, 1970; Cernusak *et al*., 2019). Even mosses with a RH_cw_ close to 100% under optimum hydric conditions (*Polytrichum commune* and *Thuidium delicatulum*) showed significant differences between Ψ_cw_ and Ψ_cyt_ under desiccation, also suggesting an active mechanism of water control but probably not constitutive or not so advanced as the rest of the studied species. Non-stomatal mechanisms for water control have been shown to be widespread in vascular plants, including C_3_ and C_4_ angiosperms (Márquez *et al*., 2024) and gymnosperms (Cernusak *et al*., 2018). Considering this generality of non-stomatal control in vascular plants, it is plausible that there is a common monophyletic origin rooted on the first land colonizers ancestors, as has also been suggested for aquaporins (Borstlap, 2002; Danielson & Johanson, 2008). In fact, mosses also seem to share the AQP-dependent biochemical mechanisms that enhance CO_2_ diffusion through the mesophyll in vascular plants (Flexas *et al*., 2006). This is evidenced by the mosses’ internal CO_2_ conductance response to temperature (Perera-Castro *et al*., 2020b) and by their apparently non-limiting cell wall thickness, been the relationship between CO_2_ assimilation rates and cell wall thickness reported for vascular plants (Flexas *et al*., 2021) elusive in mosses when big datasets are compiled for bryophytes (Perera-Castro *et al*., 2022a).

Knowing the definitive nature of this water control mechanism would shed light on its evolutionary history. A combination of cell membrane properties (e.g. aquaporin density and activity), osmotic adjustments within the apoplast, or cell wall channelling (akin to cuticular channels) especially as leaves contort, would be strong candidates for this possibly monophyletic mechanism of non-stomatal water control. Whether this mechanism is not only functionally shared across plant groups but also structurally conserved as an evolutionary pathway remains speculative. A comprehensive study, including data from ferns and lycophytes (currently unavailable to the best of our knowledge), is needed to fully support this. Nonetheless, the occurrence of a similar mechanism in both mosses and angiosperms suggests that it may also exist in these groups. Additionally, it is a priority to conclusively identify the membrane structures responsible for this mechanism in each plant group.

### Possible evolutionary constraints to water control in bryophytes

The range of RH_cw_ and its independence from the phylogeny of the studied species suggest certain interspecific plasticity of the evidenced water control in mosses. The relationships between such RH_cw_ and the pressure-volume curve derived parameters (Fig. 5) reflect a possible evolutionary constraint to the efficiency of this non-stomatal water control within mosses. In vascular plants, higher leaf capacitance has been conceived as an evolutionary advantage, since species with this characteristic are able to deal with higher and more variable evaporative demand, concomitant with the highest saturated water content, assimilation rates and lamina hydraulic conductance (Sack *et al*., 2003; Nadal *et al*., 2018; Xiong & Nadal, 2020). Shoot capacitance and elasticity are not associated with assimilation rates in bryophytes (Perera-Castro *et al*., 2020a), and *A*_opt_ and *E*_opt_ are completely independent in the studied species. Although limiting factors for CO_2_ and H_2_O diffusion seem to be unrelated in bryophytes, having dynamic storage capacity that serves as a buffer to prevent water fluctuations still can be advantageous for mosses with an “avoidance” strategy similar to those proposed by Vitt *et al*. (2014) and Jabłońska *et al*. (2023) but at shoot level. Such an avoidance strategy would be associated with higher water control and, therefore, lower RH_cw_ values.

Mosses with higher capacitance ‒ i.e. higher change in water content for a given water potential interval ‒ could maximize carbon balance by delaying the decrease in water potential. The assimilation rate, although not maximal, still can be positive in mosses for a relative water content higher than 40% (Perera-Castro *et al*., 2020a). Furthermore, low dehydration rates are necessary to guarantee the capacity to recover functionality after rehydration in some species of mosses (Schonbeck & Bewley, 1981; Proctor *et al*., 2007; Cruz de Carvalho *et al*., 2015, 2017; Yuqing *et al*., 2021) and even to prevent desiccation at all, something of vital importance in sensitive species of mosses like *Leucobryum* spp (Takács *et al*., 1999). The succulence provoked by water storage tissues with flexible cell walls has been considered a drought avoidance mechanism independent of osmotic adjustment in vascular plants (Bartlett *et al*., 2012). Since osmolyte accumulation must imply an energetic and nutritional cost in plant cells, it is consistent that the species with high water control and capacitance also presented the lowest osmotic potential.

### Conclusions

Our results indicate that mosses exhibit an active water control mechanism not previously reported. The variation in transpiration and relative humidity of the cell wall, along with the differences between cell wall and cytoplasm water potential, support the idea of a non-stomatal water control mechanism in mosses. The magnitude of the decrease in cell wall water potential correlated with traits associated with an avoidance strategy for coping with fluctuations in water availability in mosses — including cytosolic water storage, high tissue capacitance and elasticity, and a lack of osmotic adjustment. This mechanism of water loss control through cell membranes and/or cell walls is possibly derived from common ancestral traits shared with vascular plants, related to the unsaturated conditions of internal airspaces. The nature of this mechanism is unknown. Our findings reveal that mosses, due to their simple structure and the absence of stomata in the gametophyte, provide key insights into the nature and evolutionary significance of this mechanism. This makes them a valuable model for further exploration.

## ACKNOWLEDGEMENTS

AV Perera-Castro gratefully acknowledges financial support for this research by the Fulbright U.S. Student Program, which is sponsored by the U.S. Department of State and the Spanish Fulbright Commission. Its contents are solely the responsibility of the authors and do not necessarily represent the official views of the Fulbright Program, the Government of the United States, or the Spanish Fulbright Commission. DA Márquez and FA Busch were supported by the Natural Environment Research Council (grant number NE/W00674X/1). We are grateful to Prof. Becky Bixby for her help in the use of the microscope, Mr. Laura Green for technical support during the conduct of this study, and Dr. Louis Shogo and Mr. Trinity Griffus for their help in mosses collection.

## AUTHOR CONTRIBUTIONS

AVPC, FB, DM and DH contributed to the design of the study and the interpretation of the results; AVPC performed data collection, analysis, and writing of the original draft. All authors contributed to the final version of the manuscript.

## DATA AVAILABILITY

All data supporting the findings of this study are available from the corresponding author, Alicia V. Perera-Castro, upon request.

---

**Fig. S1.**
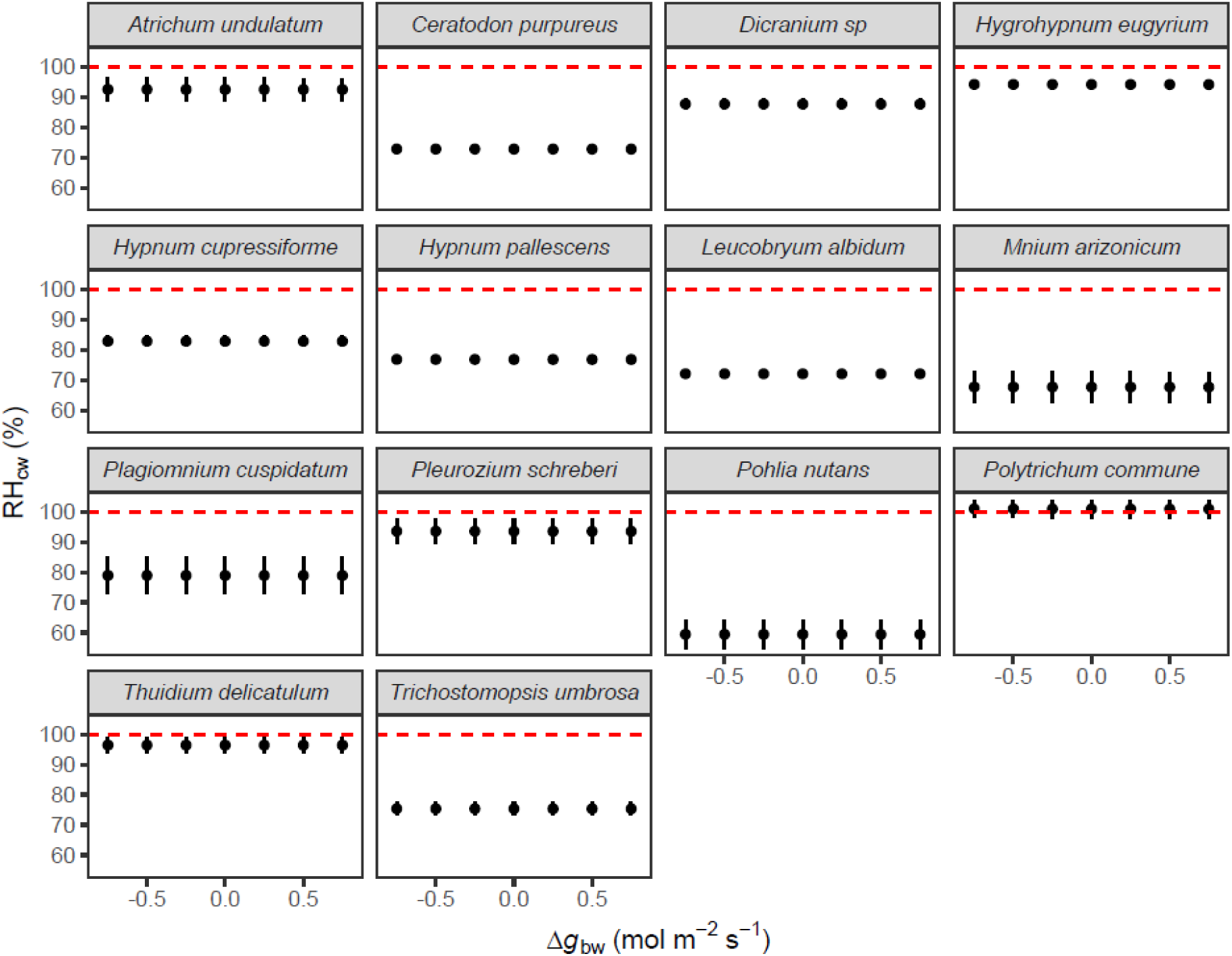
Sensitivity analysis of relative humidity of the cell wall (RH_cw_) of all studied species in response to variations of boundary layer conductance (Δ*g*_bw_) of ± 0.75 mol m^-2^ s^-1^ relative to the original recorded values (1.7-2 mol m^-2^ s^-1^). Red line indicates RH_cw_ of 100%.

**Fig. S2.**
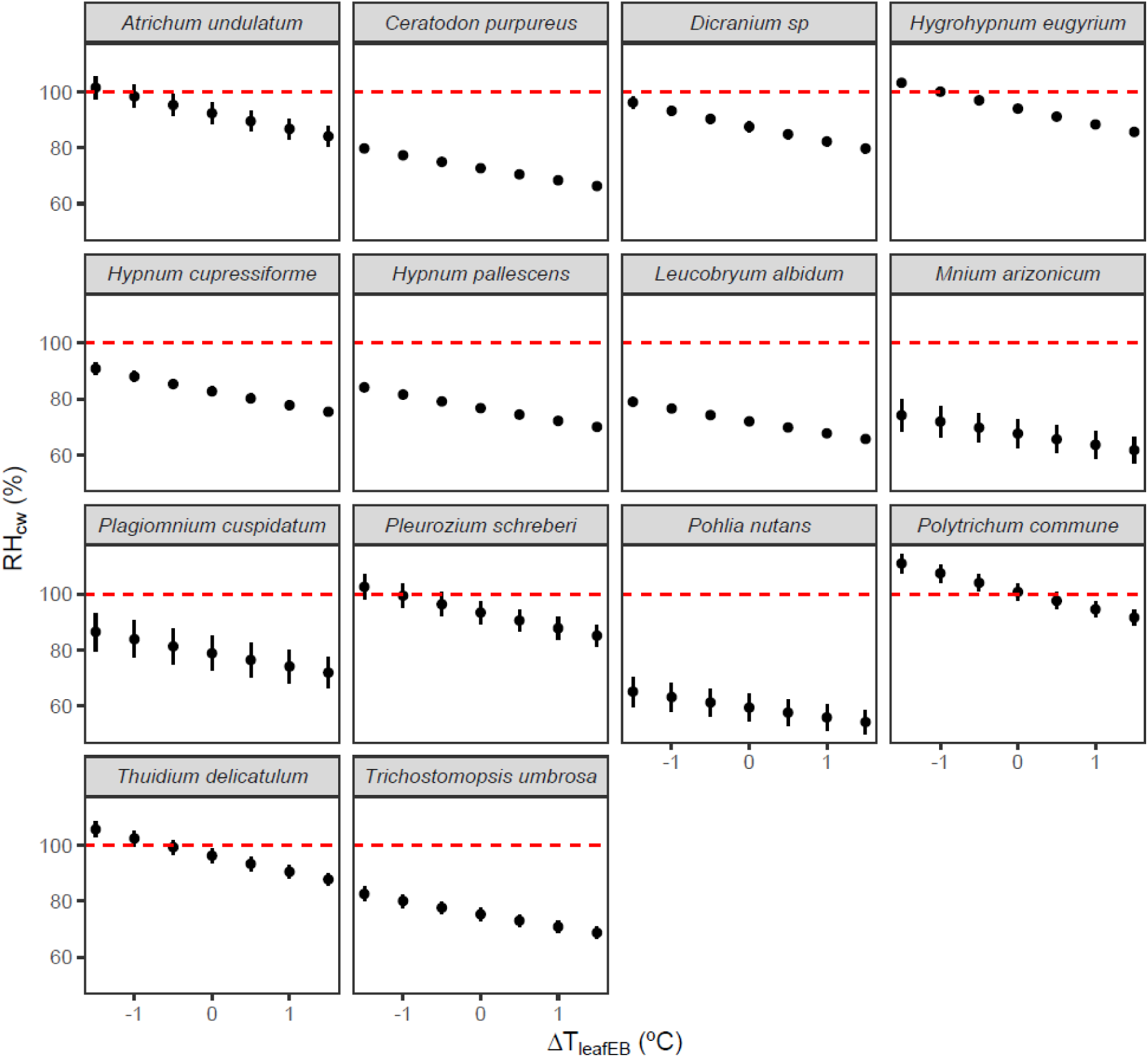
Sensitivity analysis of relative humidity of the cell wall (RH_cw_) of all studied species in response to variations of moss temperature estimated by energy balance (Δ*T*_leafEB_) of ± 1.5°C relative to the original recorded values (17.7-21.5°C). Red line indicates RH_cw_ of 100%.

**Fig. S3.**
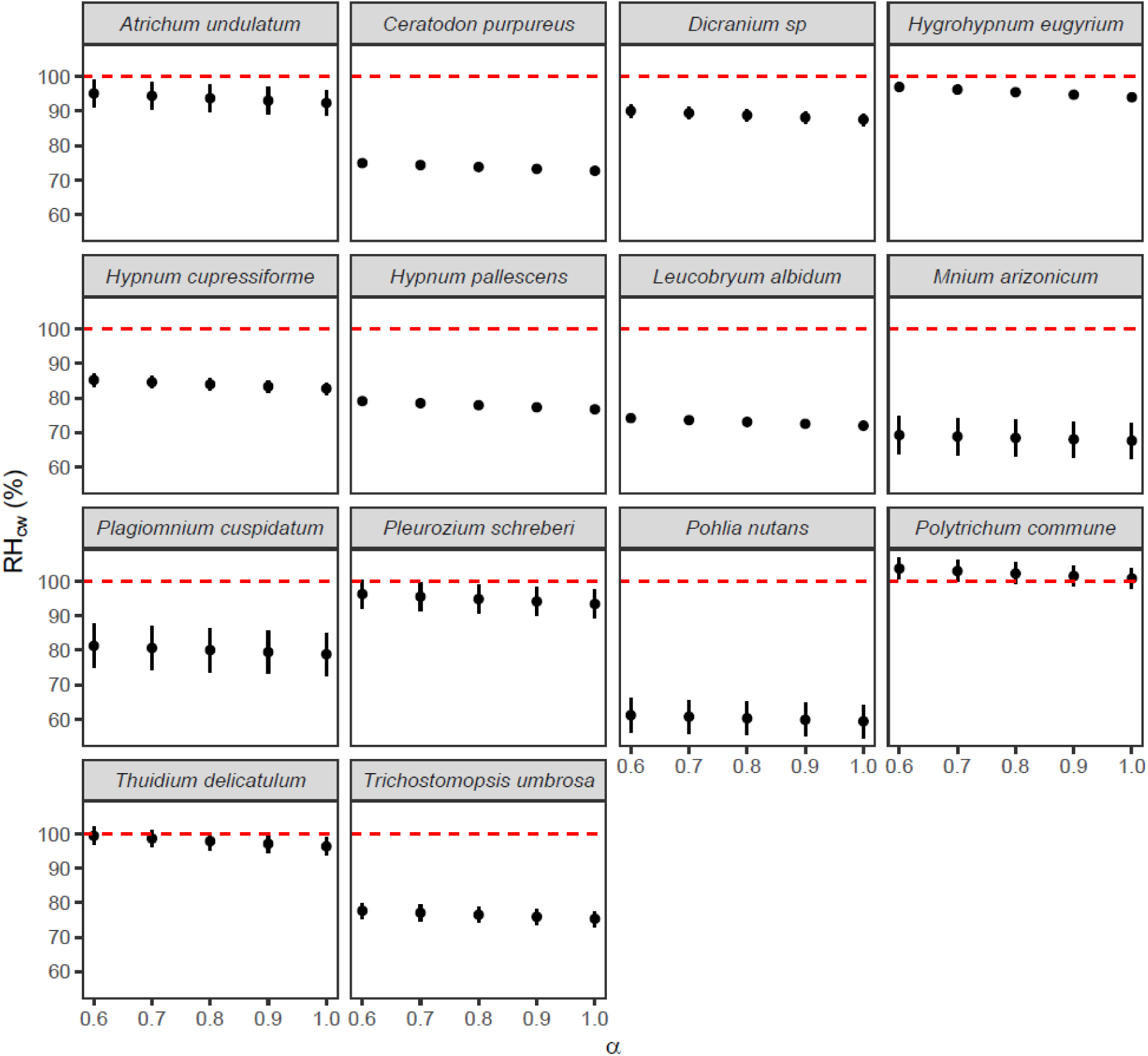
Sensitivity analysis of relative humidity of the cell wall (RH_cw_) of all studied species in response to moss absorbance of shorter wavelength (α). An ample range of α between 0.6 and 1 was assumed. Red line indicates RH_cw_ of 100%.

**Fig. S4.**
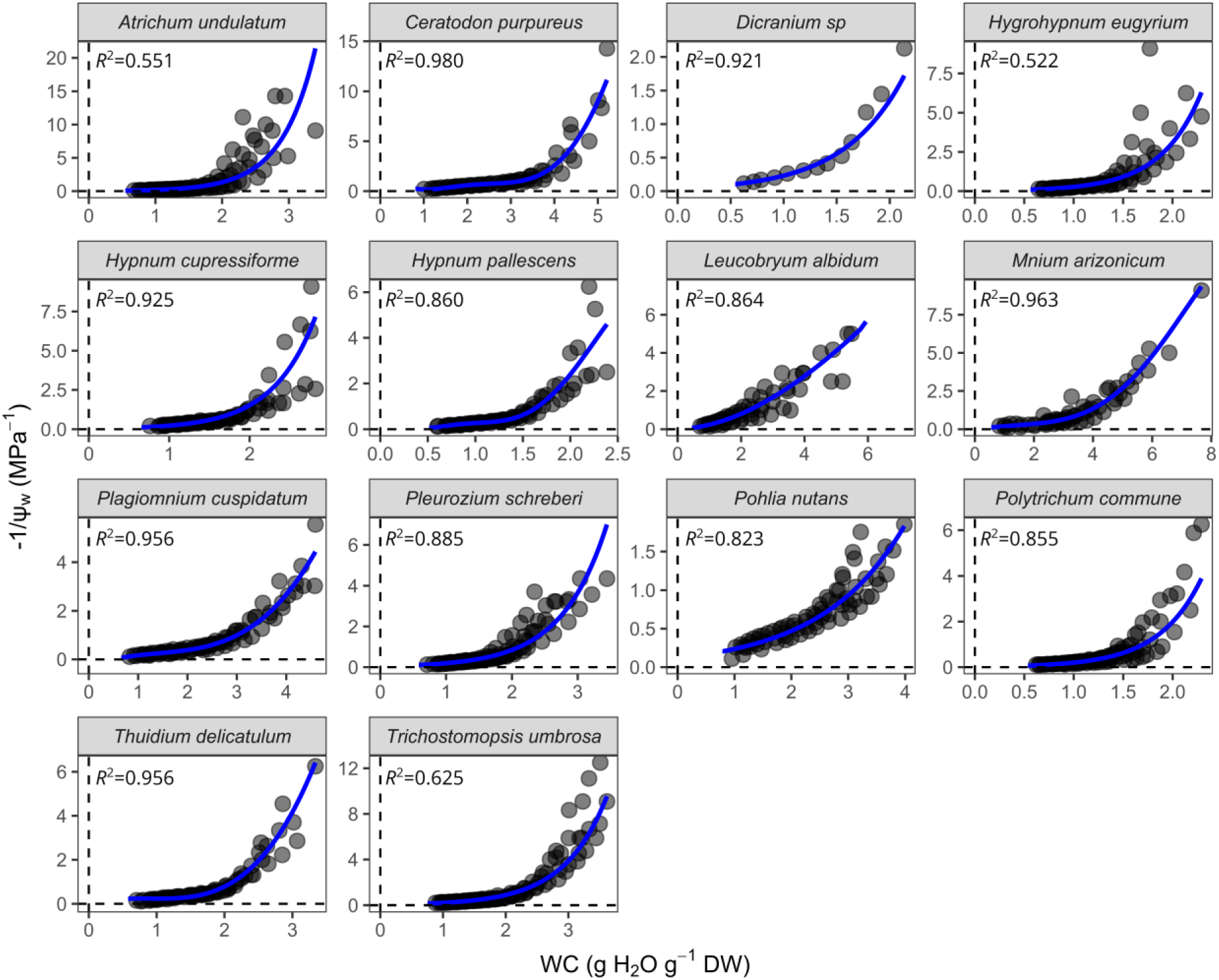
Pressure−volume curves obtained for all studied species. Blue line indicates the polynomial or exponential adjustment of the relationship between negative inverse of water potential (-1/Ψ_w_) and water content (WC).

**Fig. S5.**
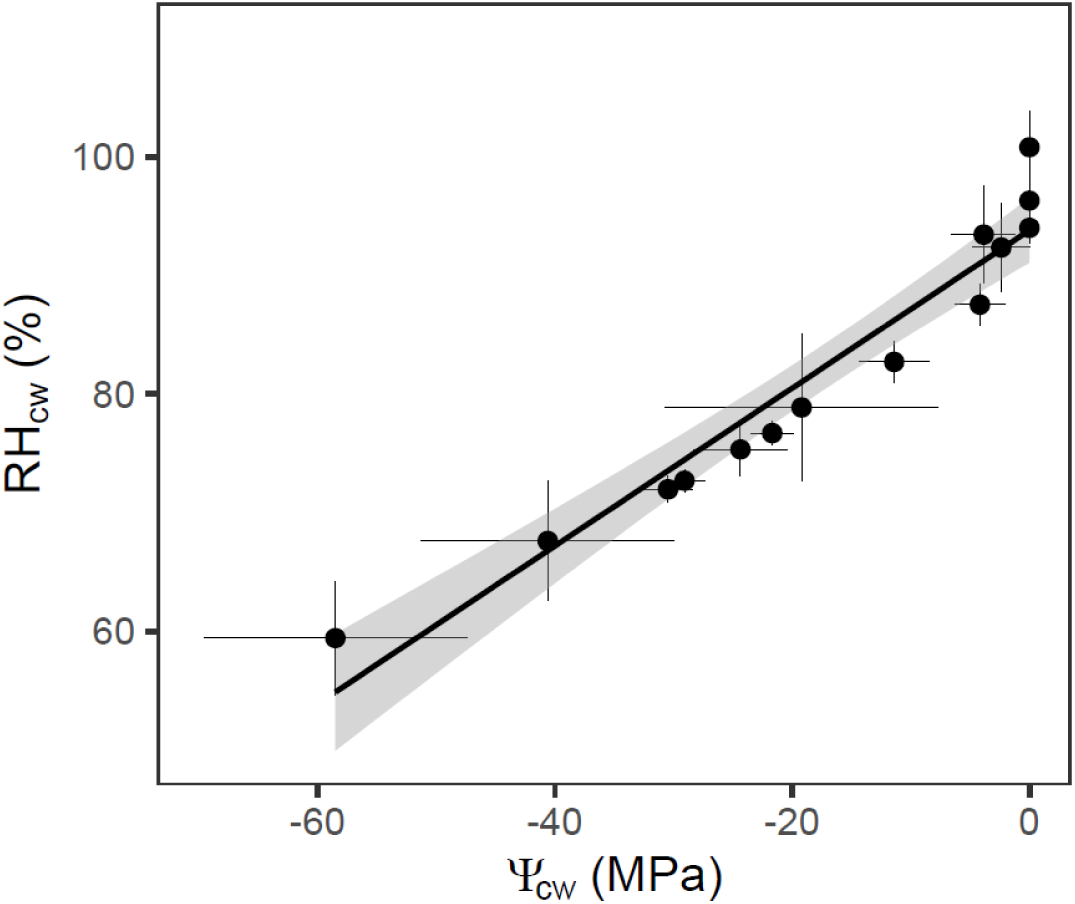
Relationship between relative humidity at cell wall (RH_cw_) at optimum water content (at maximum assimilation rate) and cell wall water vapor potential (Ψ_cw_) obtained from Eq 4 of the main manuscript. Points represent the mean ±SE for each species.

**Fig. S6.**
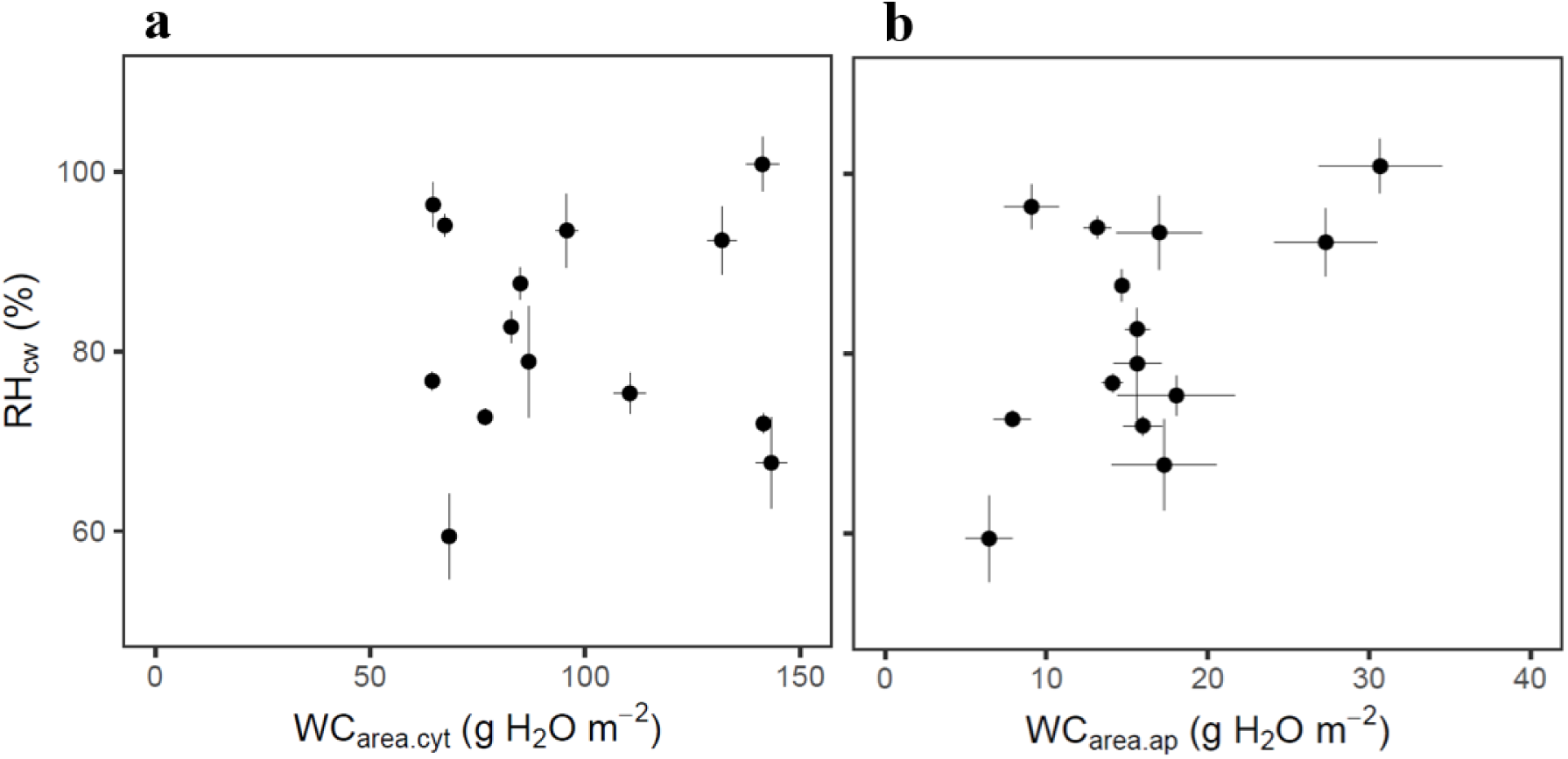
Relationship between relative humidity at cell wall (RH_cw_) at optimum water content (at maximum assimilation rate) and saturated water content of (a) cytosol and (b) apoplast per unit shoot area (WC_area.cyt_ and WC_area.ap_, respectively). Points represent the mean ±SE for each species.

